# The *C. elegans* homolog of the *Evi1* proto-oncogene, *egl-43*, coordinates G1 cell cycle arrest with pro-invasive gene expression during anchor cell invasion

**DOI:** 10.1101/802355

**Authors:** Ting Deng, Michael Daube, Alex Hajnal, Evelyn Lattmann

## Abstract

Cell invasion allows cells to migrate across compartment boundaries formed by basement membranes. Aberrant cell invasion is a first step during the formation of metastases by malignant cancer cells.

Anchor cell (AC) invasion in *C. elegans* is an excellent *in vivo* model to study the regulation of cell invasion during development. Here, we have examined the function of *egl-43*, the homolog of the human *Evi1* proto-oncogene (also called *MECOM*), in the invading AC. *egl-43* plays a dual role in this process, firstly by imposing a G1 cell cycle arrest to prevent AC proliferation, and secondly, by activating pro-invasive gene expression. We have identified the AP-1 transcription factor *fos-1* and the *Notch* homolog *lin-12* as critical *egl-43* targets. A positive feedback loop between *fos-1* and *egl-43* induces pro-invasive gene expression in the AC, while repression of *lin-12 Notch* expression by *egl-43* ensures the G1 cell cycle arrest necessary for invasion. Reducing *lin-12* levels in *egl-43* depleted animals restored the G1 arrest, while hyperactivation of *lin-12* signaling in the differentiated AC was sufficient to induce proliferation.

Taken together, our data have identified *egl-43 Evi1* as a critical factor coordinating cell invasion with cell cycle arrest.

**Author summary:** Cells invasion is a fundamental biological process that allows cells to cross compartment boundaries and migrate to new locations. Aberrant cell invasion is a first step during the formation of metastases by malignant cancer cells.

We have investigated how a specialized cell in the Nematode *C. elegans*, the so-called anchor cell, can invade into an adjacent epithelium. Our work has identified an oncogenic transcription factor that controls the expression of specific target genes necessary for cell invasion, and at the same time inhibits the proliferation of the invading anchor cell.

These findings shed light on the mechanisms, by which cells decide whether to proliferate or invade.

## Introduction

Cell invasion, which is initiated by the breaching of basement membranes (BMs), is a regulated physiological process allowing selected cells to cross compartment boundaries during normal development. Cell invasion is also the first step that is activated during the formation of metastases by malignant cancer cells [1]. Anchor cell (AC) invasion in *C. elegans* is a genetically amenable and trackable model that has provided important insights into the molecular pathways regulating cell invasion and uncovered the molecular similarities between tumor cell and developmental cell invasion [2,3].

The AC is specified during the second larval stage (L2) of *C. elegans* development, when two equivalent precursor cells (Z1.ppp and Z4.aaa) adopt either the AC or the ventral uterine (VU) fate, depending on stochastic differences in LAG-2 Delta/ LIN-12 Notch signaling [4,5]. The cell that exhibits higher *lag-2* expression levels adopts the “default” AC fate and activates LIN-12 Notch signaling in the adjacent cell to induce the VU fate [6]. The AC never divides, while the VU cell undergoes three rounds of cell divisions. AC invasion occurs during the third larval stage (L3) of *C. elegans* development, in order to form a direct connection between the uterus and the vulval epithelium [7]. In this process, the AC breaches two BM layers and establishes direct contact with the invaginating vulval epithelium.

Specification of the VU fate depends on the activity of the e*gl-43* gene, which encodes two isoforms of a Zinc finger transcription factor homologous to the mammalian *Evi1* proto-oncogene in the MECOM locus [8–10]. Human *Evi1* has been implicated in the development of different types of cancer, most notably in the hematopoietic system in acute myeloid leukemia (AML) [11]. Inhibition of *C. elegans egl-43* during the mid-L2 stage leads to the formation of two ACs due to a defect in VU cell specification [8]. However, *egl-43* remains expressed in the AC after its specification, and EGl-43 is required together with the AP-1 transcription factor FOS-1 in the invading AC to induce the expression of genes that promote BM breaching (i.e. pro-invasive genes), such as *zmp-1*, *cdh-3* and *him-4* [8,9,12]. AC invasion also depends on the transcription factor NHR-67, a nuclear receptor of the *tailless* family, which maintains the G1 arrest of the AC [13].

Despite the importance of *egl-43* in activating the expression of pro-invasive genes and BM breaching, its exact role during AC invasion remains poorly understood. Here, we report that *egl-43* and *fos-1* form a positive auto-regulatory feedback-loop that drives pro-invasive gene expression in the AC. Moreover, *egl-43* plays a previously unknown role in establishing the G1 arrest of the AC by repressing Notch-dependent AC proliferation. Thus, *egl-43* acts as an important regulator of AC invasion that coordinates the G1 arrest with pro-invasive gene expression.

## Results

### Deletion of the long *egl-43L* isoform leads to the formation of multiple ACs that fail to invade

In order to study the role of the different *egl-43* isoforms during AC invasion, we used CRISPR/Cas9 genome editing [14] to insert *gfp* tags at the 5’ or 3’ end of the *egl-43* locus (Fig. 1A). Since the transcriptional start site of the short isoform (*egl-43S*) is located within the 5th intron of the *egl-43L* locus, the insertion of the *gfp* cassette at the 5’ end of the first exon labels exclusively the protein encoded by the long isoform (*gfp::egl-43L*), whereas the insertion of the *gfp* tag at the 3’ end of the locus labels both, the short *egl-43S* and the long *egl-43L* isoforms (*egl-43::gfp*, Fig. 1A). Furthermore, the *gfp* cassette inserted at the 5’ end contained two FRT sites in the *gfp* introns, permitting the tissue-specific inactivation of the *egl-43L* isoform (suppl. Fig S1). For the expression analysis, we focused on the mid-L3 stage (the Pn.pxx stage, after the VPCs had undergone two rounds of cell divisions), the time when AC invasion normally occurs [7]. Both reporters, *gfp::egl-43L* and *egl-43::gfp*, were expressed in the invading AC and in the surrounding ventral and dorsal uterine (VU and DU) cells (Fig. 1B, rows 1 and 2). We found no obvious difference in the expression pattern of the two *egl-43* reporters, suggesting that the long *egl-43L* isoform accounts for most of the expression observed. In order to test if the uterine expression is indeed caused by *egl-43L*, we designed two RNAi clones, one specifically targeting *egl-43L* (exons 4 and 5 of *egl-43L*), and the other targeting the 5’ UTR of *egl-43S,* which is not included in the *egl-43L* transcript (Fig. 1A). EGL-43::GFP expression was lost upon *egl-43L* RNAi, yet no difference in expression was observed after *egl-43S* RNAi (Fig. 1C), supporting the above conclusion that the uterine expression originates predominantly from the long *egl-43L* isoform.

**Figure 1.**
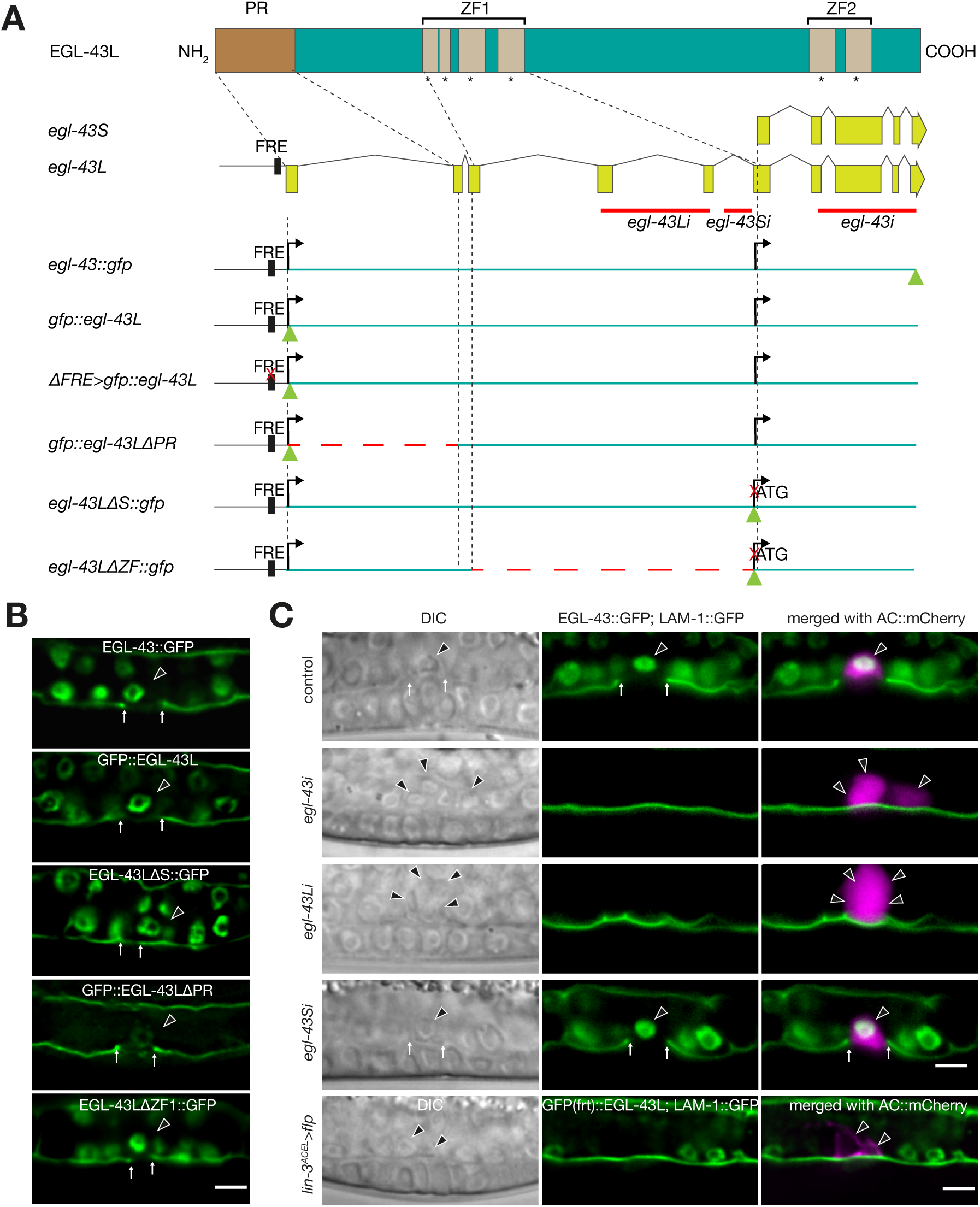
Loss of *egl-43L* function leads to the formation of multiple ACs. **(A)** Schematic overview of the *egl-43* locus and the CRISPR/Cas9 engineered alleles used in this study. Green triangles indicate *gfp* insertion sites and dashed red line indicate deleted regions. The red crosses indicate the sites of the 11 bp (TTACTCATCTT) deletion in the Fos-Responsive Element (FRE) and of the mutation in the initiation codon of the *egl-43S* isoform. Solid red lines refer to the regions targeted by RNAi. (**B**) Expression patterns of the different endogenous *egl-43* reporters depicted in (**A**). The basement membrane (BM) was simultaneously labelled with the LAM-1::GFP marker to score AC invasion. **(C)** RNAi of the different *egl-43* isoforms and FLP/FRT-mediated excision of *egl-43L*. Left panels show Nomarski (DIC) images and middle panels the fluorescence signals of endogenous EGL-43::GFP expression in the nuclei and the LAM-1::GFP BM marker. The right panels show the GFP signals merged with the ACs labelled by the *lin-3^ACEL^>mCherry* reporter (rows 1-4) and *cdh-*3>mCherry::PH (row 5) in magenta. The black arrowheads point at the AC nuclei and the white arrows at the locations of the BM breaches. Control refers to animals exposed to the empty RNAi vector. The scale bars are 5 µm.

As reported previously [9], *egl-43L* RNAi led to an invasion defect with a penetrance comparable to that of total *egl-43* RNAi (91% for *egl-43L* (n=34) and 95% for total *egl-43* RNAi (n=22)). Consistent with a role of *egl-43* during AC/VU specification [8,9], we detected early L3 larvae with two ACs upon *egl-43L* or total *egl-43* RNAi (Fig. 1C). However, in 17 out of 22 cases, total *egl-43* RNAi led to the formation of more than two ACs, a phenotype that cannot be explained by an AC/VU specification defect (Fig. 1C). Similar to the invasion defects, the occurrence of multiple ACs was also caused by selective inhibition of the long *egl-43L* isoform, with 26 out of 34 worms containing more than 2 ACs (Fig. 1C). Moreover, the number of ACs increased progressively with the age of the larvae, indicating an ongoing proliferation of the AC after the L2 stage (Fig. 2A, B). This pointed at an additional role of *egl-43L* in preventing the proliferation of the AC after its specification, independently of its function during VU fate specification.

**Figure 2.**
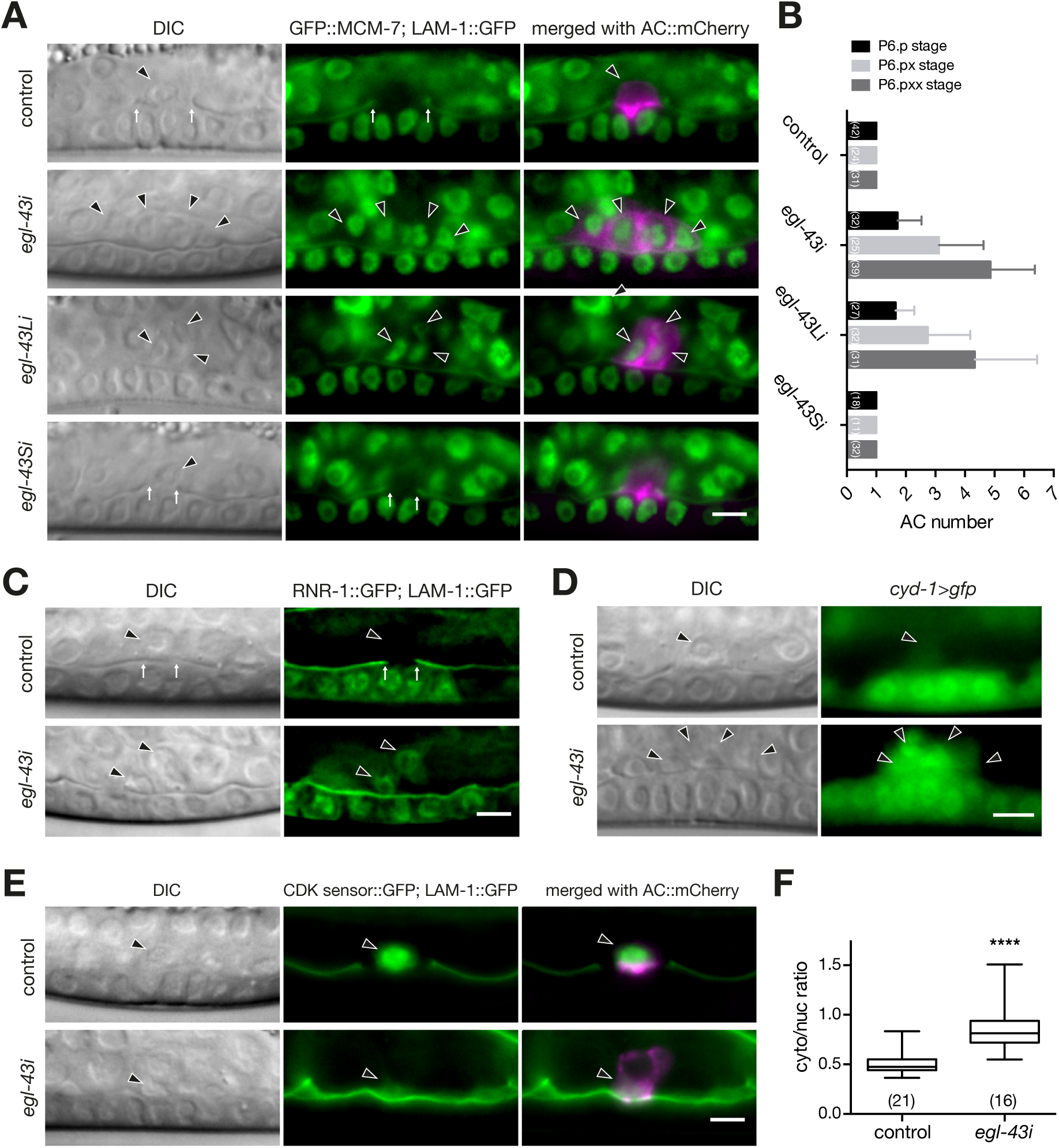
*egl-43* is required for the G1 arrest of the AC. **(A)** Expression of the endogenous GFP::MCM-7 reporter after RNAi of the different *egl-43* isoforms. The left panels depict Nomarski (DIC) images and the middle panel nuclear GFP::MCM-7 expression together with the LAM-1::GFP BM marker at the mid-L3 stage. The right panels show the GFP signals merged with the ACs labelled by the *cdh-3*>mCherry::PH reporter in magenta. **(B)** Quantification of the number of ACs formed after RNAi of the different *egl-43* isoforms from the early L3 (Pn.p) until the mid-L3 (Pn.pxx) stage. The error bars indicate the standard deviation. **(C)** Expression of the S-phase marker RNR-1::GFP together with LAM-1::GFP in control and *egl-43* RNAi-treated mid-L3 larvae. **(D)** Expression of the CyclinD *cyd-1>gfp* reporter in control and *egl-43* RNAi-treated mid-L3 larvae. **(E)** Expression of the CDK activity sensor in the AC of control and *egl-43* RNAi-treated mid-L3 larvae together with the LAM-1::GFP BM marker. The right panels show the CDK sensor signal merged with the ACs labelled by the *lin-3^ACEL^>mCherry* reporter in magenta. **(F)** Quantification of CDK sensor activity in the AC. Box plots showing the cytoplasmic to nuclear intensity ratio as a measure of CDK activity. The error bars indicate the Min-to-Max values. Statistical significance was determined with a Student’s t-test and is indicated with **** for p<0.0001. In each graph, the numbers of animals scored are indicated in brackets. The black arrowheads point at the AC nuclei and the white arrows at the locations of the BM breaches. The scale bars are 5 µm.

In order to specifically examine the role of the *egl-43S* isoform during AC invasion, we used CRISPR/Cas9 engineering to introduce an ATGàCTG mutation in the predicted *egl-43S* start codon (*egl-43LΔS::gfp,* Fig. 1A, B). The *egl-43LΔS::gfp* strain was viable and showed a similar expression pattern as the *egl-43::gfp* and *gfp::egl-43L* strains (Fig. 1B, rows 1-3). Moreover, AC invasion occurred normally in all *egl-43LΔS::gfp* animals examined, and no AC proliferation was observed (n=35), which confirms the predominant role of *egl-43L* in regulating AC invasion and proliferation.

Finally, we used the Flp/FRT system to generate an AC-specific knock-out of *egl-43L* [15]. Since two FRT sites had been inserted in introns of the *gfp* cassette at the 5’ end of the *egl-43L* locus, the expression of the FLPase under control of the AC-specific *lin-3* enhancer element (*lin-3^ACEL^>flp*) [16] specifically inactivated *egl-43L* in the differentiated AC (suppl. Fig. S1). Deletion of *egl-43L* in the AC led to invasion defects (9 out of 17 animals) and the formation of multiple ACs (4 out of 17 cases), similar to the phenotypes caused by *egl-43* RNAi (Fig. 1C, **row 5)**. Thus, *egl-43L* acts cell-autonomously in the AC to prevent proliferation and promote invasion.

### Neither the N-terminal PR nor the ZF1 domains in EGL-43 are necessary for AC invasion

*egl-43L* encodes a transcription factor containing an N-terminal PRD1-BF1/RIZ-1 domain (PR) and two separate clusters of Zinc finger domain (ZF1 & ZF2) that could exhibit DNA binding activity [10]. PR domains are structurally similar to the SET (Su(var)3-9, Enhancer of zeste and Trithorax) domains, which contribute to histone lysine methyltransferase activity and have also been reported to mediate protein-protein interactions [17,18]. In order to dissect the requirement of the different EGL-43 domains during AC invasion, we generated in-frame deletions in the *egl-43* coding region using the CRISPR/Cas9 system (Fig. 1A). In the PR domain deletion allele (*gfp::egl-43ΔPR*, deleted amino acids 2-62), we detected an approximately 75% decrease in GFP::EGL-43L expression levels in the AC (Fig. 1B, row 4; suppl. Fig S2A). Despite this strong reduction, neither the proliferation nor the invasion of the AC were affected (all 25 animals scored showed normal AC invasion and no proliferation).

Thus, the PR domain in EGL-43 is not necessary for AC invasion. Though, the reduced expression levels suggested that the PR domain is either required for *egl-43* autoregulation (see below), protein stability, or that there exist additional regulatory elements in the deleted first intron (Fig. 1A) that promote *egl-43* expression in the AC.

No AC invasion defects or change in expression levels were observed in the *egl-43ΔZF1::gfp* mutant, which is an in frame deletion removing amino acids 161-235 containing the N-terminal Zinc Finger domains (ZF1) (Fig. 1B, row 5; suppl. Fig S2B). By contrast, the *egl-43* null mutant *tm1802*, which contains a 659 bp deletion removing the C-terminal Zinc Finger domains (ZF2), displays severe developmental defects and early embryonic lethality [9]. Thus, the regulation of AC invasion and proliferation most likely depends on the C-terminal ZF2 domains.

### *egl-43* is required for the G1 arrest of the AC

Previous studies have shown that the AC must remain arrested in the G1 phase in order to invade [13]. The occurrence of multiple (i.e. more than two) ACs upon inactivation of *egl-43L* indicated that the AC had bypassed the G1 arrest and begun to proliferate. In order to test this hypothesis, we performed RNAi knock down of the different *egl-43* isoforms in a strain carrying the cell cycle marker GFP::MCM-7 [19,20]. *mcm-7* encodes a subunit of the pre-replication complex (pre-RC) that is highly expressed in cycling cells but down-regulated in non-proliferating cells. No GFP::MCM-7 expression was detectable in the AC after control or *egl-43S* RNAi. However, *egl-43* and *egl-43L* RNAi resulted in elevated GFP::MCM-7 levels in the multiple ACs that formed, indicating that these ACs had re-entered the cell cycle (Fig. 2A). Consistent with this notion, the average number of ACs increased during the development from the Pn.p (late L2/early L3) stage to the Pn.pxx (mid to late L3) stage (Fig. 2B).

To further characterize the cell cycle state of the AC, we analyzed RNR-1::GFP expression, which serves as an S-phase marker [20]. While RNR-1::GFP was absent in the AC of control RNAi animals, it was expressed in the multiple ACs formed after *egl-43* RNAi (Fig. 2C). Moreover, we detected elevated levels of a transcriptional *cyd-1>gfp* Cyclin D reporter in the multiple ACs produced after *egl-43i*, while the single AC in control animals showed only faint Cyclin D expression (Fig. 2D). Finally, we expressed a newly developed CDK biosensor in the AC to directly quantify CDK kinase activity [21]. This sensor monitors CDK activity via the phosphorylation-induced nuclear export of a GFP-tagged kinase substrate. Thus, a high cytoplasmic-to-nuclear signal ratio indicates high CDK activity. The multiple ACs formed in *egl-43* RNAi-treated animals showed a significantly increased CDK sensor activity compared to the single AC in control RNAi animals (Fig. 2E, F).

Thus, loss of EGL-43 function increases CDK activity and triggers cell cycle entry of the AC.

### *egl-43* acts independently of *nhr-67* and *hda-1* to induce the G1 arrest of the AC

As reported previously [13], RNAi of *nhr-67* led to the appearance of multiple ACs that could not breach the BM (Fig. 3A). Since loss of *nhr-67* led to a similar phenotype as inhibition of *egl-43*, we tested whether the expression of *nhr-67* depends on *egl-43*, or vice versa. In order to avoid artefacts caused by the dilution of the signal in the multiple, smaller ACs of mid-L3 larvae, we quantified reporter expression in early L3 larvae at the Pn.p stage, before AC proliferation occurred. Quantification of endogenous GFP::EGL-43L reporter levels after *nhr-67* RNAi revealed no change in *egl-43* expression (Fig. 3A, B). Conversely, expression of an *nhr-67::gfp* reporter was not significantly changed by *egl-43* RNAi (Fig. 3C, D). The histone deacetylase gene *hda-1* acts downstream of *nhr-67* to induce the G1 arrest of the AC [13]. However, the expression levels of an HDA-1::RFP reporter in the AC were not changed by *egl-43* RNAi (Fig. 3E, F). These data suggested that *egl-43* is expressed independently of *nhr-67* and *hda-1* to cause the cell cycle arrest of the AC.

**Figure 3.**
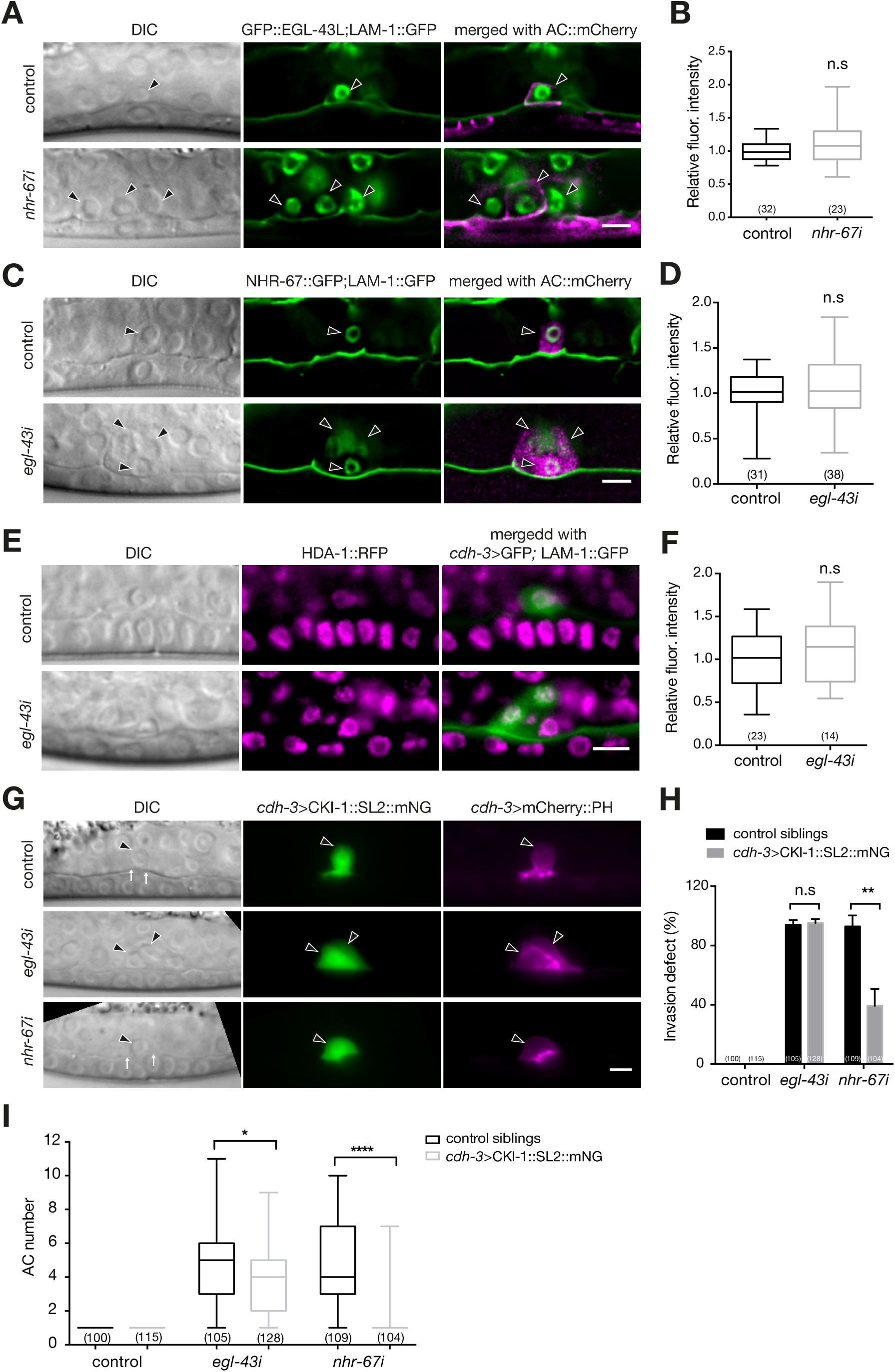
*egl-43* acts independently of *nhr-67* and *hda-1*. **(A)** Expression of GFP::EGL-43L after control and *nhr-67* RNAi. The left panels depict Nomarski (DIC) images and the middle panels GFP::EGL-43L expression together with the LAM-1::GFP BM marker at the early-L3 (Pn.p) stage. The right panels show the GFP signals merged with the ACs labelled by the *cdh-3>mCherry::PH* reporter in magenta. **(B)** Quantification of GFP::EGL-43L levels in the AC. **(C)** Expression of NHR-67::GFP after control and *egl-43* RNAi. The left panels depict Nomarski (DIC) images and the middle panels NHR-67::GFP expression together with the LAM-1::GFP BM marker at the early-L3 (Pn.p) stage. The right panels show the GFP signals merged with the ACs labelled by the *lin-3^ACEL^>mCherry* reporter in magenta. **(D)** Quantification of NHR-67::GFP levels in the AC. **(E)** Expression of HDA-1::RFP after control and *egl-43* RNAi. The left panels depict Nomarski (DIC) images and the middle panels HDA-1::RFP expression in magenta at the mid-L3 stage. The right panels show the RFP signal merged with the ACs labelled by the *cdh-3*>GFP marker together with LAM-1::GFP in green. **(F)** Quantification of HDA-1::RFP levels in the AC. **(G)** AC-specific expression of *cki-1* from the *cdh-3* enhancer/promoter in control, *egl-43* and *nhr-67* RNAi-treated mid-L3 larvae. The left panels depict Nomarski (DIC) images, the middle panels CKI-1::SL2::mNG expression in green and the right panels the ACs labelled by the *cdh-3>*mCherry::PH reporter in magenta. **(H)** Quantification of the AC invasion and **(I)** proliferation phenotypes in RNAi-treated animals expressing CKI-1::SL2::mNG (gray bars) compared to their control siblings lacking the *cdh-3>cki-1::SL2::mNG* transgene (black bars/boxes). The error bars in the box plots in **(B)**, **(D)**, **(F)** and **(I)** indicate the Min-to-Max values, and the error bars in the bar chart in **(H)** indicate the standard deviation. Statistical significance was determined with a Student’s t-test and is indicated with n.s. for p>0.05, * for p<0.05, ** for p<0.01, and **** for p<0.0001. The black arrowheads point at the AC nuclei. The numbers in brackets in the graphs refer to the numbers of animals analyzed. The scale bars are 5 µm.

Due to the high penetrance of the AC proliferation phenotype caused by *nhr-67* or *egl-43* RNAi alone, we were unable to test if the simultaneous knock-down of both transcription factors had an additive effect. Consistent with a previous report [13], the AC-specific expression of the CDK inhibitor *cki-1* (together with an mNeonGreen (mNG) marker on a bi-cistronic mRNA) under control of a *cdh-3* enhancer/promoter fragment (*cdh-3>cki-1::SL2::mNG*) rescued the AC proliferation and invasion defects caused by *nhr-67* RNAi (Fig. 3G, H). By contrast, CKI-1 overexpression did not rescue the invasion defects caused by *egl-43* knockdown, and it only partially blocked AC proliferation (Fig. 3G, H, I).

These data indicated that *egl-43* and *nhr-67* maintain the cell cycle arrest of the AC through different mechanisms. We thus postulated that *egl-43* acts independently of *nhr-67* and *hda-1* to promote the G1 arrest and invasion of the AC.

### *egl-43* inhibits AC proliferation by repressing *lin-12 Notch* expression

LIN-12 Notch signaling is not only critical during the AC/VU decision, but it also links cell differentiation to cell cycle progression in different other tissues [4,22,23]. We therefore tested whether *egl-43* regulates *lin-12 Notch* expression in the AC by examining a translational LIN-12::GFP reporter [24]. In control animals at the mid-L3 stage, LIN-12::GFP expression had disappeared in the AC, but expression persisted in the adjacent VU cells (Fig. 4A). By contrast, the multiple ACs that formed after *egl-43* RNAi continued to express LIN-12::GFP (Fig. 4A). The ACs in *egl-43* RNAi-treated larvae still expressed a *lin-3::mNG* reporter, which serves as a marker distinguishing the AC from the VU fate (suppl. Fig. S3A) [25], as well as a reporter for the LIN-12 ligand LAG-2 (suppl. Fig. S3B) [26]. Thus, the inhibition of *egl-43* did not cause a transformation of the AC into a VU fate, but rather resulted in the ectopic expression of LIN-12 in the proliferating ACs.

**Figure 4.**
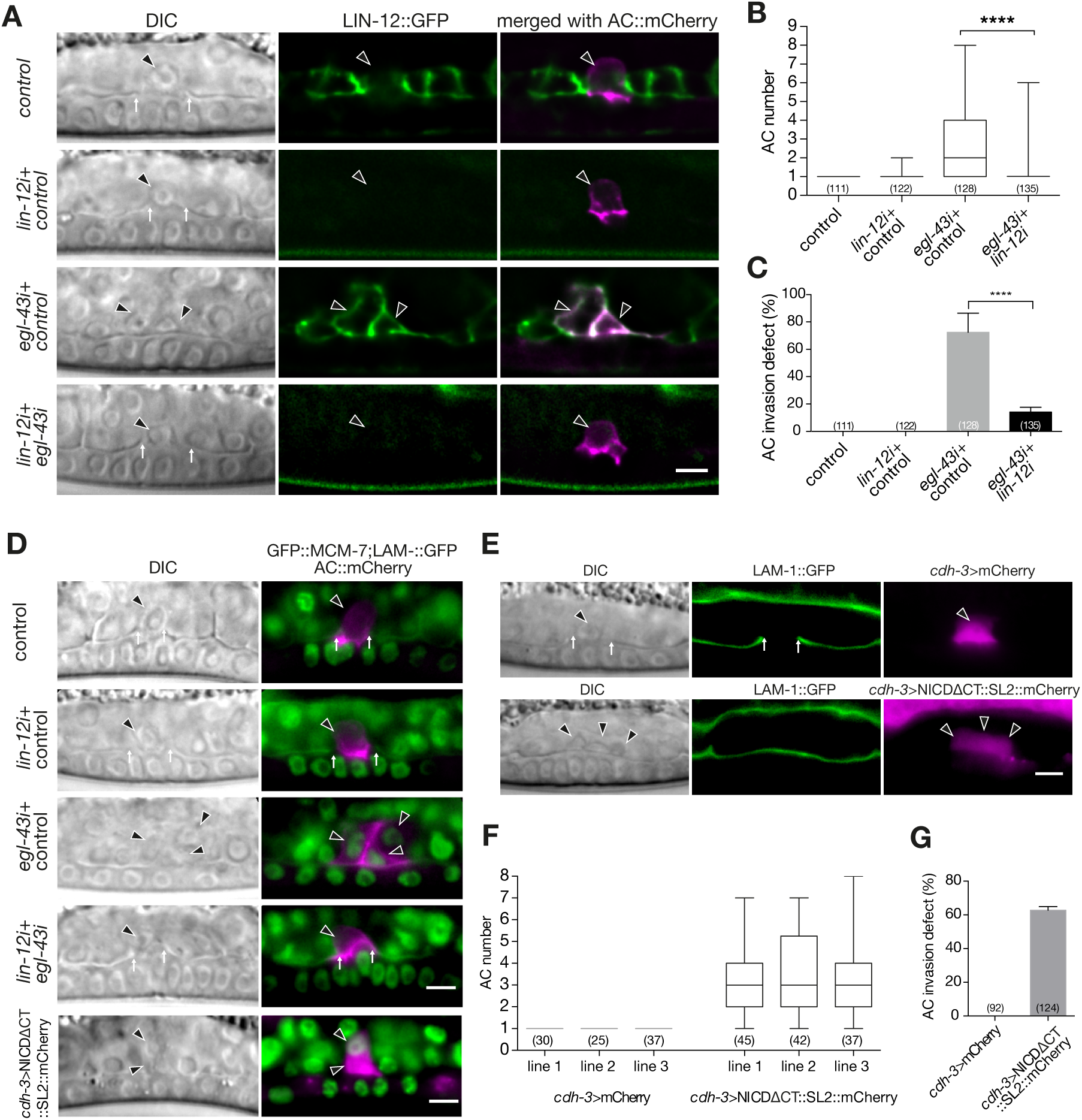
*egl-43* represses *lin-12 Notch* expression to prevent AC proliferation. **(A)** Expression of LIN-12::GFP after control, *egl-43* or *lin-12* single and *egl-43; lin-12* double RNAi. The left panels shows Nomarski (DIC) images, middle panels LIN-12::GFP expression in green and the right panels the GFP signal merged with the AC labelled by the *cdh-3>mCherry::PH* reporter in magenta **(B)** Quantification of the AC numbers in mid-L3 larvae treated with the different RNAi combinations shown in **(A)**. **(C)** Quantification of the AC invasion defects caused by the different RNAi treatments shown in **(A)**. **(D)** GFP::MCM-7 expression together with LAM-1::GFP after control, *egl-43* or *lin-12* single and *egl-43; lin-12* double RNAi, and in a larva expressing NICDΔCT::mCherry. The left panels show Nomarski (DIC) images and the right panels the GFP::MCM-7 signal in green merged with the AC labelled by the *cdh-3>mCherry::PH* reporter in magenta. **(E)** AC-specific expression of *nicdΔct* from the *cdh-3* enhancer/promoter leads to the formation of multiple ACs. Left panels shows Nomarski (DIC) images and middle panels the BM marker LAM-1::GFP. The right panels show the mCherry expression from the *cdh-3* promoter as a control (row 1) or co-expressed with *nicdΔct* from a bi-cistronic transcript (row 2). **(F)** Quantification of the AC numbers in three independent control lines expressing mCherry alone and in three lines expressing NICDΔCT together with mCherry in the AC. The error bars in **(B)** & **(F)** indicate the Min-to-Max values and in **(C)** and **(G)** the standard deviation. Statistical significance was determined with a Student’s t-test and is indicated with **** for p<0.0001. The black arrowheads point at the AC nuclei. The numbers in brackets in the graphs refer to the numbers of animals analyzed. The scale bars are 5 µm.

To test whether an ectopic activation of *lin-12* Notch signaling in the AC is responsible for the AC proliferation phenotype, we performed double RNAi of *egl-43* and *lin-12* and scored the number of ACs formed, as well as their ability to invade. While 63% of *egl-43* single RNAi-treated animals formed multiple ACs, only 16% of *egl-43*; *lin-12* double RNAi-treated animals contained multiple ACs. The average number of ACs per animals decreased from 2.6 in *egl-43* single to 1.2 in *egl-43; lin-12* double RNAi-treated animals (Fig. 4B). Also, the penetrance of the invasion defects decreased from 72% to 14% (Fig. 4C).

To test if reducing *lin-12* activity restored the cell cycle arrest of the AC, we examined the GFP::MCM-7 reporter, which is expressed exclusively in proliferating cells. Control or single *lin-12* RNAi did not induce GFP::MCM-7 expression in the AC of mid-L3 larvae, while *egl-43* RNAi resulted in the formation of multiple GFP::MCM-7 positive ACs (Figs. 2A & 4D). By contrast, the single ACs formed in *egl-43; lin-12* double RNAi-treated animals did not express GFP::MCM-7 (Fig. 4D). Thus, reducing *lin-12* activity suppressed the AC multiplication and invasion defects caused by *egl-43* RNAi and restored the G1 arrest of the AC.

### Activation of LIN-12 Notch signaling in the differentiated AC triggers proliferation

These results led to the hypothesis that the ectopic activation of LIN-12 Notch signaling caused by loss of *egl-43* function may trigger the re-entry of the AC into the cell cycle. In order to test if LIN-12 signaling is sufficient to induce AC proliferation, we expressed a fragment of the intracellular LIN-12 Notch domain, in which the C-terminal PEST degradation motif had been deleted (NICDΔCT) [22], under the control of the AC-specific *cdh-3* enhancer/promoter fragment. Expression of NICDΔCT in the VPCs was shown to hyper-activate the Notch signaling pathway [22], and *cdh-3*-driven expression occurs only after the AC fate has been specified [27]. Three independent transgenic lines carrying the *cdh-3>nicdΔct::SL2::mCherry* transgene (co-expressing an mCherry marker on a bi-cistronic mRNA) exhibited an AC proliferation phenotype with an average of 3.4 ACs per animal, which is comparable to the phenotype observed after *egl-43* RNAi (Fig. 4E, F). In addition, the GFP::MCM-7 reporter was up-regulated (Fig. 4D, row 5) while GFP::EGL-43L continued to be expressed in the multiple ACs of *cdh-3>nicdΔct::SL2::mCherry* animals (suppl. Fig. S3C). Moreover, NICDΔCT expression caused a penetrant AC invasion defect (Fig. 4G).

Thus, the ectopic activation of the LIN-12 Notch pathway in the differentiated AC was sufficient to trigger cell cycle entry. Taken together, we propose that EGL-43 inhibits LIN-12 Notch expression to maintain the G1 arrest of the invading AC.

### A positive regulatory feedback loop between *egl-43* and *fos-1* activates pro-invasive gene expression

Since we found that blocking AC proliferation by overexpressing CKI-1 was not sufficient to rescue the invasion defects caused by *egl-43* RNAi (Fig. 3), we speculated that EGL-43 might perform two functions in the AC; first to repress AC proliferation and second, to activate pro-invasive gene expression. Along these lines, we have previously reported that *fos-1* positively regulates *egl-43* expression in the AC and that *egl-43* is required for the expression of *zmp-1*, *cdh-3* and *him-4* [9].

To further characterize the role of *egl-43* in regulating pro-invasive gene expression, we investigated a mutual regulation of *fos-1* and *egl-43*. The expression of the endogenous GFP::EGL-43L reporter in the AC was reduced approximately two-fold in homozygous *fos-1(ar105)* mutants compared to heterozygous *fos-1(ar105)/+* control siblings at the mid-L3 stage (Fig. 5A, B). Using CRISPR/Cas9 genome editing, we deleted the 11 bp (TTACTCATCTT) FOS-Responsive Element (FRE) [9] in the promoter region of the endogenous *gfp::egl-43L* reporter strain (*ΔFRE>gfp::egl-43L*, Fig. 1A). In a heterozygous *fos-1(ar105)/+* background, the expression of the mutant *ΔFRE>gfp::egl-43L* reporter was reduced to a similar extent as the wild-type *gfp::egl-43L* reporter was reduced in a homozygous *fos-1(ar105)* background. Since the levels of the FRE mutant reporter did not further decrease in homozygous *fos-1(ar105)* larvae, FOS-1 appears to control *egl-43* expression mainly through this FRE (Fig. 5A, B). While this experiment confirmed that FOS-1 up-regulates endogenous *egl-43* expression, it also pointed at the existence of additional factors that activate *egl-43* expression in the AC. In particular, the deletion of the FRE in *egl-43* did not cause an obvious defect in AC invasion (all 23 animals scored showed normal AC invasion), suggesting that the reduction in *egl-43* expression after the deletion of the FRE can be compensated by the AC.

**Figure 5.**
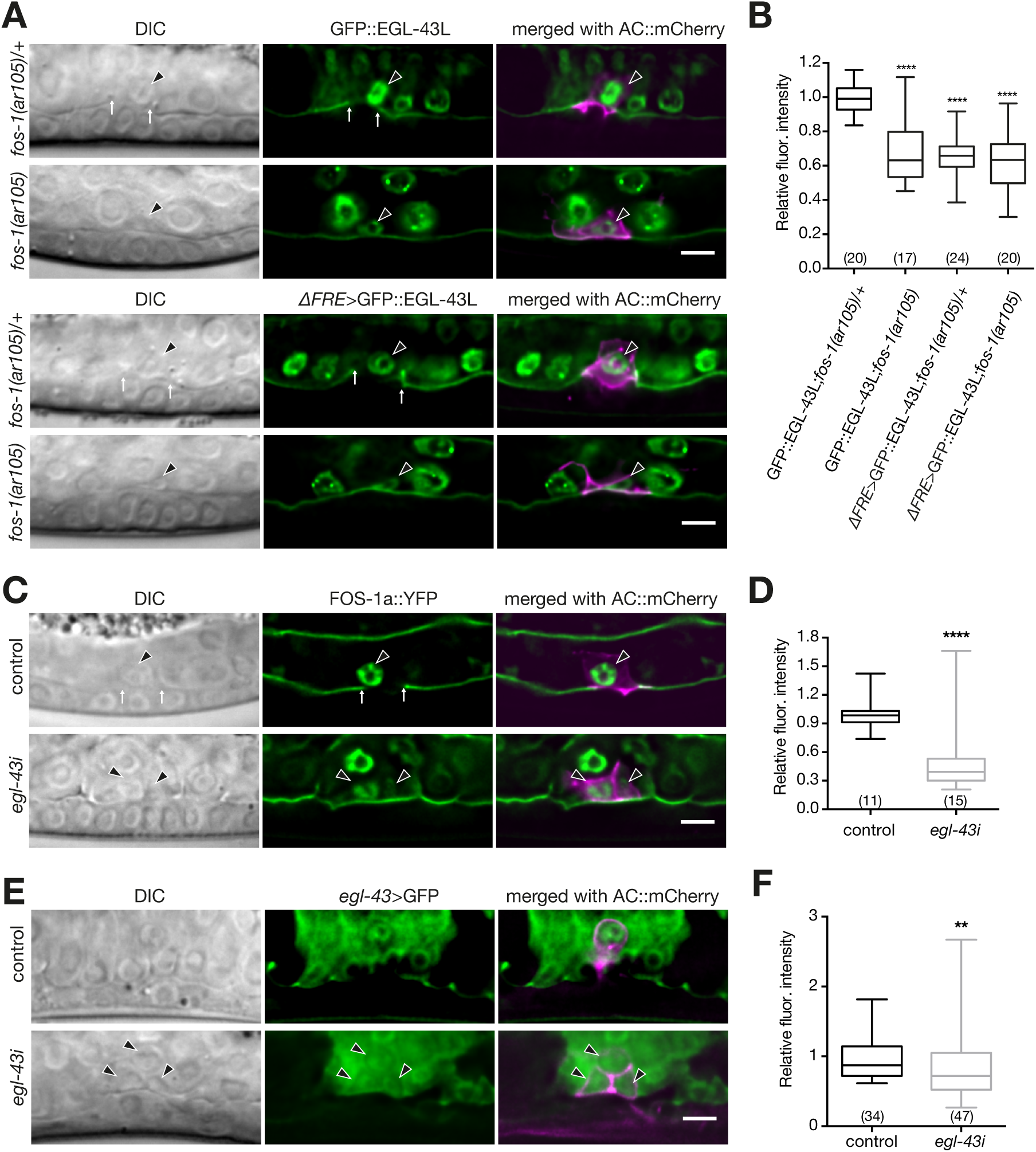
*egl-43* and *fos-1* regulate each other’s expression. **(A)** Expression of endogenous GFP::EGL-43L and of the mutant *ΔFRE*>GFP::EGL-43L reporter carrying a deletion of the Fos-responsive element TTACTCATCTT (ΔFRE), each in a *fos-1(ar105)/+* heterozygous and *fos-1(ar105)* homozygous background at the mid-L3 stage. **(B)** Quantification of GFP::EGL-43L expression levels in the ACs of the indicated mutant backgrounds. **(C)** Expression of a FOS-1a::YFP reporter after control and *egl-43* RNAi. **(D)** Quantification of FOS-1a::YFP levels in the ACs after control and *egl-43* RNAi. **(E)** Expression of a transcriptional *egl-43>gfp* reporter in the ACs after control and *egl-43* RNAi. **(F)** Quantification of a transcriptional *egl-43>gfp* reporter after control and *egl-43* RNAi. For each reporter, the left panels show Nomarski (DIC) images, the middle panels the GFP signals of the indicated reporters in green (in **(A)** and **(C)** together with the LAM-1::GFP BM marker) and the right panels the GFP reporter signals merged with the ACs labelled with the *cdh-3>mCherry::PH* reporter in magenta. The black arrowheads point at the AC nuclei and the white arrows at the locations of the BM breaches. The error bars in the box plots in **(B)**, **(D)** & **(F)** indicate the Min-to-Max values. Statistical significance was determined with a Student’s t-test and is indicated with ** for p<0.01 and **** for p<0.0001. The numbers in brackets refer to the numbers of animals analyzed. The scale bars are 5 µm.

Since *egl-43* and *fos-1* regulate some of the same target genes [9,12], we tested if *egl-43* regulates *fos-1a* expression. The expression of a *fos-1a::yfp* reporter in the AC was reduced approximately three-fold by *egl-43* RNAi (Fig. 5C, D), indicating that *egl-43* and *fos-1* positively regulate each other’s expression. Moreover, the expression of a transcriptional *egl-43L* reporter (*egl-43L>gfp*, a strain containing an insertion of the self-excising cassette [14] after the *gfp* coding sequences to terminate transcription 5’ of the *egl-43* coding sequences) was reduced in the multiple ACs that formed after *egl-43* RNAi (Fig. 5E, F). Thus, *egl-43* positively regulates its own expression in the AC.

In summary, the expression analysis indicated that *egl-43* and *fos-1* form a positive feedback loop that maintains high expression of both transcription factors in the AC to induce the expression of target genes required for invasion. The auto-regulation of *egl-43* may add further robustness to this network.

## Discussion

Uncontrolled activation of cell invasion is one of the hallmarks of malignant cancer cells that form metastases [1]. Genetic studies in model organisms have indicated that invasive tumor cells re-activate the same molecular programs that control cell invasion during normal animal development. AC invasion in *C. elegans* has served as an excellent model to dissect the genetic pathways regulating the invasive phenotype of a single cell [3].

Here, we report that the *egl-43* gene, which encodes the *C. elegans* ortholog of the human *Evi1* proto-oncogene, plays two distinct roles in the invading AC (Fig. 6). First, EGL-43 maintains the AC arrested in the G1 phase of the cell cycle by repressing the expression of the LIN-12 Notch receptor. The ectopic activation of Notch signaling in the differentiated AC was sufficient to induce proliferation in the presence of EGL-43 expression, indicating that LIN-12 acts downstream of EGL-43 to reactivate the cell cycle in the AC. Likewise, a recent study in *C. elegans* has highlighted the importance of the LIN-12 Notch pathway in keeping an equilibrium between the proliferation and differentiation of somatic cells [23].

**Figure 6.**
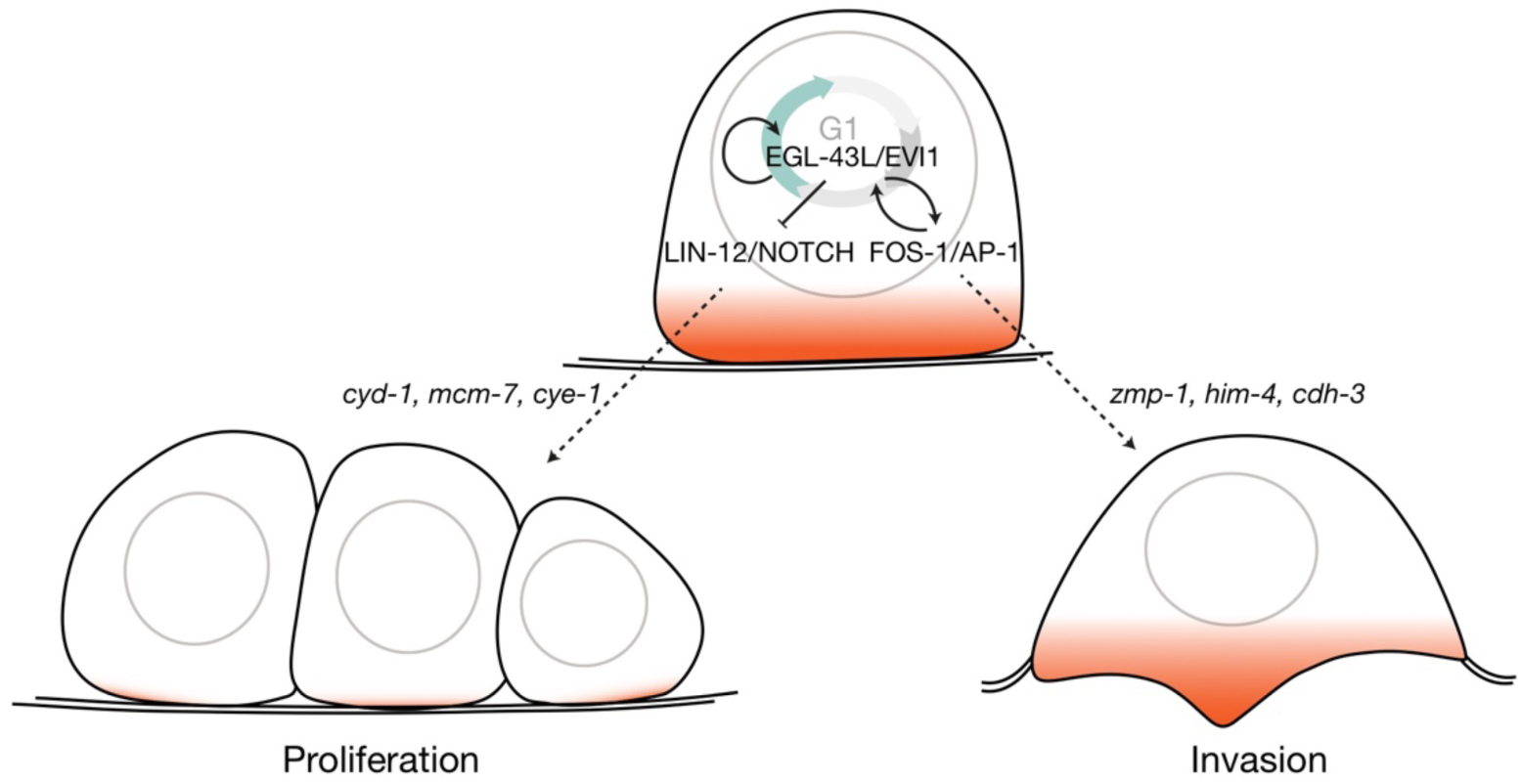
Dual function of EGL-43L during AC invasion. EGL-43L has two distinct functions during AC invasion. Left side: EGL-43L represses LIN-12 Notch expression in the differentiated AC to maintain the G1 arrest and prevent proliferation. Right side: EGL-43L activates in a positive feedback loop together with FOS-1 the expression of pro-invasive genes in the AC.

It was previously shown that the nuclear receptor NHR-67 is required for the G1 arrest of the AC and that the cell cycle arrest is a prerequisite for AC invasion [13]. However, our data indicate that EGL-43 and NHR-67 act in parallel through different mechanisms, since blocking the cell cycle of the AC by overexpressing the CDK inhibitor CKI-1 rescued the AC invasion defects caused by *nhr-67* RNAi, but not those caused by *egl-43* RNAi. Hence, EGL-43 appears to perform additional functions in the AC besides merely inhibiting cell cycle progression.

Our data indicate that EGL-43 is part of a regulatory network controlling the expression of pro-invasive genes in the AC. We have previously identified *egl-43* as a downstream target of *fos-1* based on transgenic reporter analysis [9]. By deleting the FOS-1 responsive element in the endogenous *egl-43* locus, we could confirm that *egl-43L* expression is indeed positively regulated by FOS-1. However, this experiment also indicated that the regulation of *egl-43L* by FOS-1 is not essential for AC invasion. Hence, additional factors must exist that induce *egl-43* expression in parallel with FOS-1. Furthermore, EGL-43 activates *fos-1* as well as its own expression in the AC. Interestingly, a similar auto-activating activity has been described for mammalian *Evi1* in hematopoietic stem cells [10]. Thanks to these positive feedback loops between *egl-43* and *fos-1*, low levels of either of the two transcription factors may be sufficient to induce a stable expression of both transcription factors and thereby irreversibly determine the invasive fate of the AC.

Taken together, EGL-43 coordinates the expression of pro-invasive genes with the cell cycle state of the AC by activating *fos-1* and at the same time repressing *lin-12 Notch* expression. Thus, EGL-43 is a central component in a regulatory network, which decides whether cells divide or invade.

## Material and Methods

### C. elegans culture and maintenance

*C. elegans* strains were maintained at 20°C on standard nematode growth plates as described [28]. The wild-type strain was *C. elegans* Bristol, variety N2. We refer to translational protein fusions with a :: symbol between the gene name and the tag, while transcriptional fusions are indicated with a > symbol between the promoter/enhancer and the tag. The genotypes of the strains used in this study are listed in suppl. Table S1. The construction of the plasmids, oligonucleotides and sgRNAs used to generate the different reporters are described in the **Extended Methods** and in suppl. Tables S2-S4.

### Scoring the AC invasion phenotype

AC invasion was scored in mid-L3 larvae after the VPC had divided twice (Pn.pxx stage) as described [9]. We monitored the continuity of the BM by DIC or fluorescence microscopy using the *qyIs10*[*lam-1>lam-1::gfp*] transgene as a marker.

### RNA interference

RNAi interference was done by feeding dsRNA-producing *E. coli* [29]. Larvae were synchronized at the L1 stage by hypochlorite treatment of gravid adults and plated on NGM plates containing 3mM IPTG. P0 animals were analyzed after 30-36 hours of treatment. For double RNAi experiments, bacteria of the indicated clones were mixed at a 1:1 ration. RNAi clones targeting genes of interest were obtained from the *C. elegans* genome-wide RNAi library or the *C. elegans* open reading frame (ORFeome) RNAi library (both from Source BioScience). RNAi vectors targeting *egl-43L* and *egl-43S* were subcloned into the L4440 vector by Gibson assembly (suppl. Table S3). The empty L4440 vector (labelled “control” in the figures) was used as negative control in all experiments.

### Microscopy and image analysis

Fluorescent and Nomarski images were acquired with a LEICA DM6000B microscope equipped with a Leica DFC360 FX camera and a 63x (N.A. 1.32) oil-immersion lens, or with an Olympus BX61 wide-field microscope equipped with a X-light spinning disc confocal system, a 100x Plan Apo (N.A. 1.4) lens and an iXon ultra888 EMCCD camera. Worms were imaged with 100 x magnification and z-stacks with a spacing of 0.1–0.8 μm were recorded. The Fiji software [30] was used for image analysis and fluorescent intensity quantifications using the built-in measurement tools. Figures were assembled with Adobe Illustrator software.

## Acknowledgements

We wish to thank members of the Hajnal laboratory for numerous discussions, the *Caenorhabditis Genetics Center*, which is funded by NIH Office of Research Infrastructure Programs (P40 OD010440), and the van der Heuvel laboratory for providing strains. We are also grateful to Andrew Fire for making *gfp* vectors available, and to Julie Ahringer for providing RNAi clones. The Hajnal laboratory is supported by the Swiss National Science Foundation grant no. 31003A-166580 and by the Kanton of Zürich.

## Supplementary information

**Figure S1.**
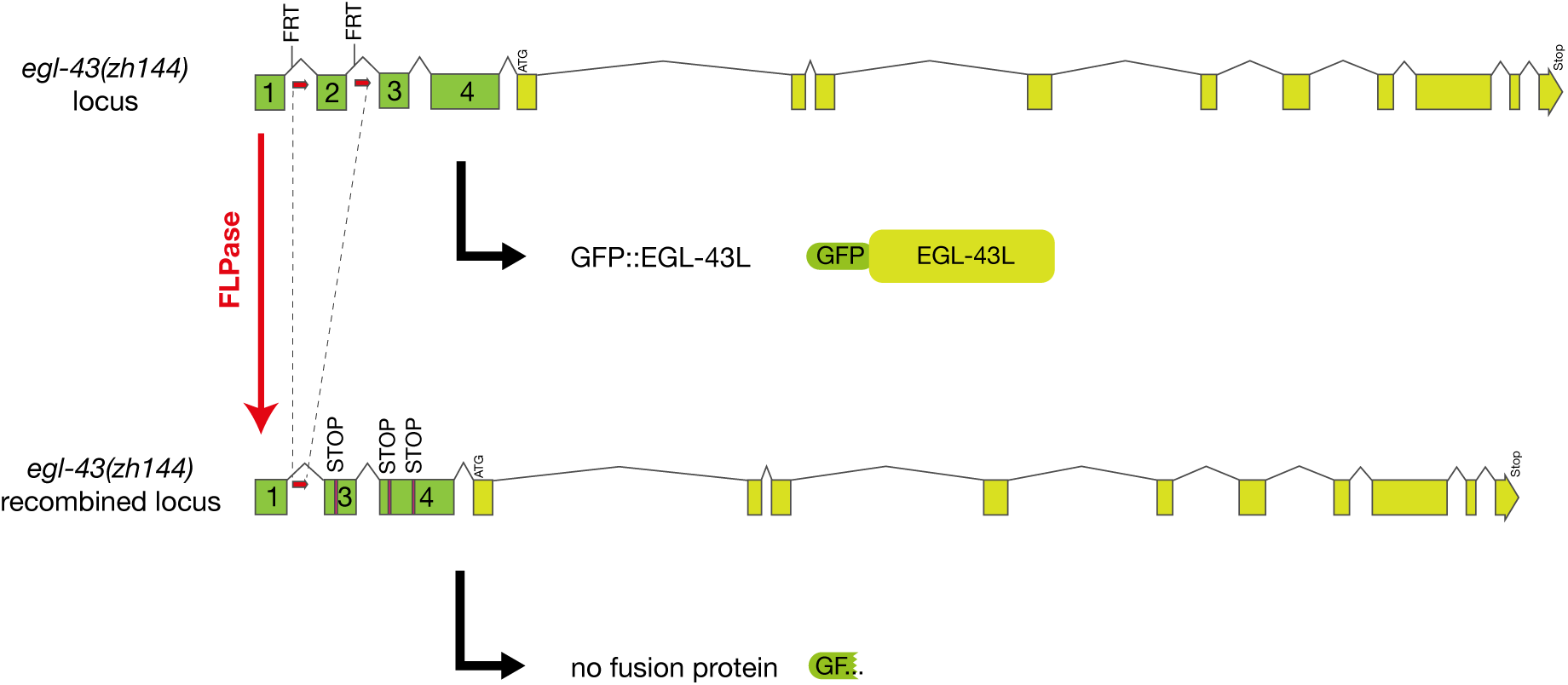
Structure of the FRT-tagged *gfp::egl-43L* allele *zh144*.

**Figure S2.**
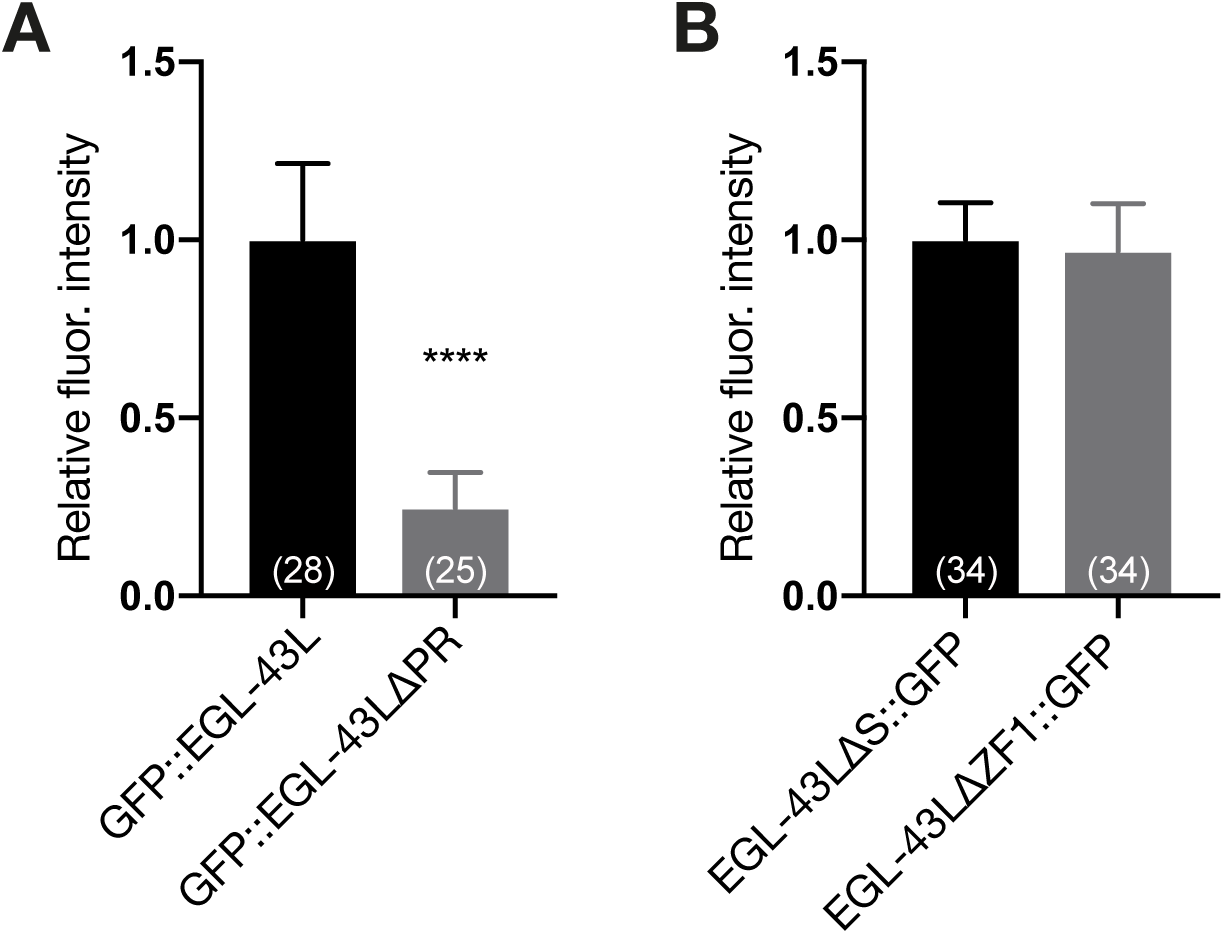
The PR domain deletion reduces *egl-43* expression levels. **(A)** Quantification of GFP::EGL-43L and GFP::EGL-43LΔPR and (**B**) EGL-43LΔS::GFP and EGL-43LΔZF1::GFP expression levels in mid-L3 larvae.

**Figure S3.**
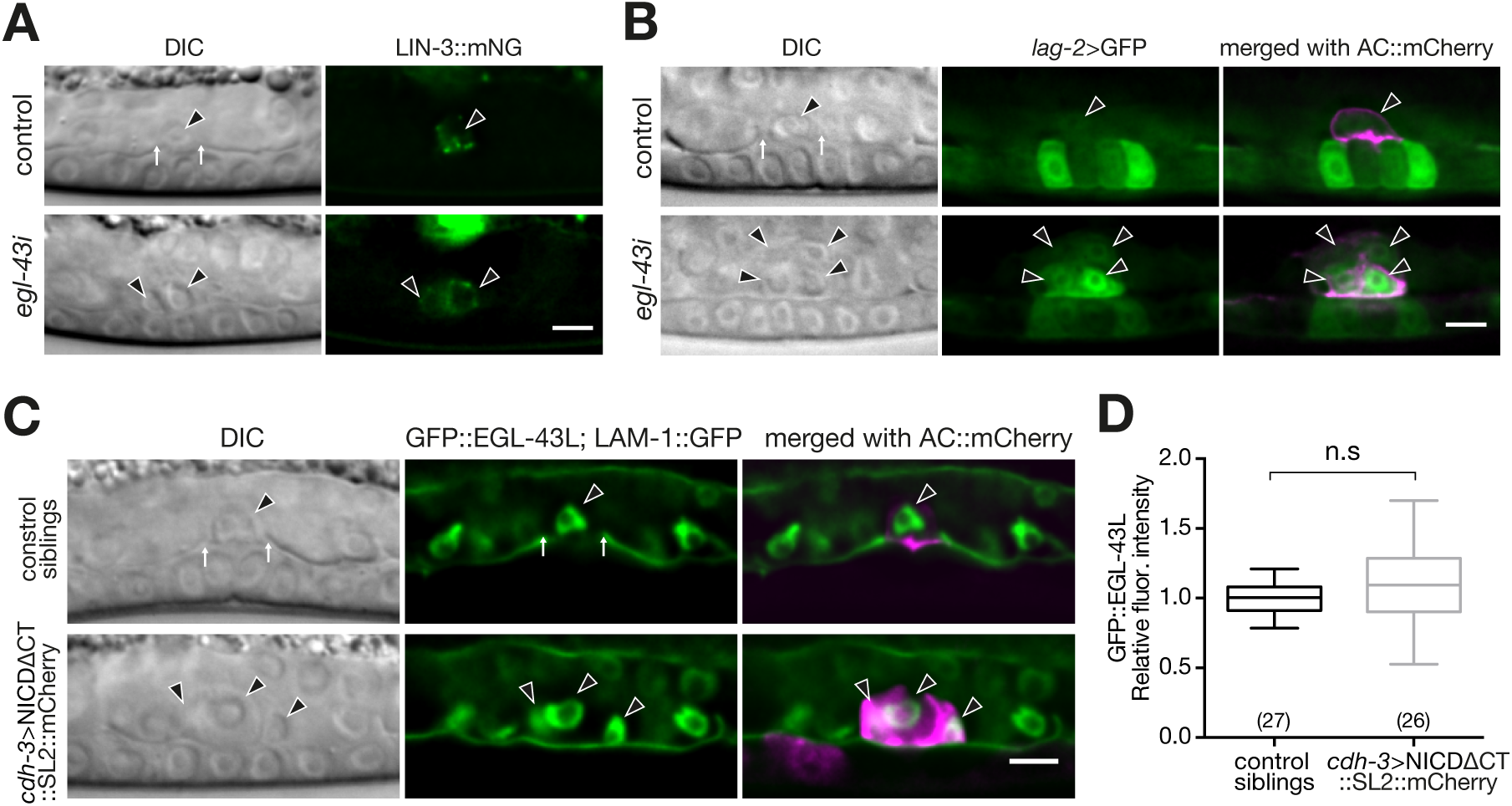
AC fate markers remain expressed after *egl-43* RNAi or Notch activation. **(A)** *lin-3* reporter expression in the control and *egl-43* RNAi ACs. Left panels show Nomarski (DIC) images and right panels the fluorescence image with LIN-3::mNG. **(B)** *lag-2* expression in the ACs of control and *egl-43* RNAi. Left panels shows Nomarski (DIC) images, middle panel the fluorescence image with *lag-2*>GFP reporter, and right panels the reporter merged with AC marker *cdh-3*>mCherry::PH. **(C)** GFP::EGL-43L expression in the ACs of control and NICDΔCT expressing ACs. Left panels show Nomarski (DIC) images, middle panels the GFP::EGL-43 signal with the LAM-1::GFP BM marker, and right panels merged with the ACs labelled with *cdh-3*>PH::mCherry (control, row 1) and *cdh-3*>NICDΔCT::SL2::mCherry (row 2) respectively. **(D)** Quantification of C. Two-tail student test was used. n.s indicates p>0.05 (p=0.0627). The scale bars are 5 µm.

**Table S1.**
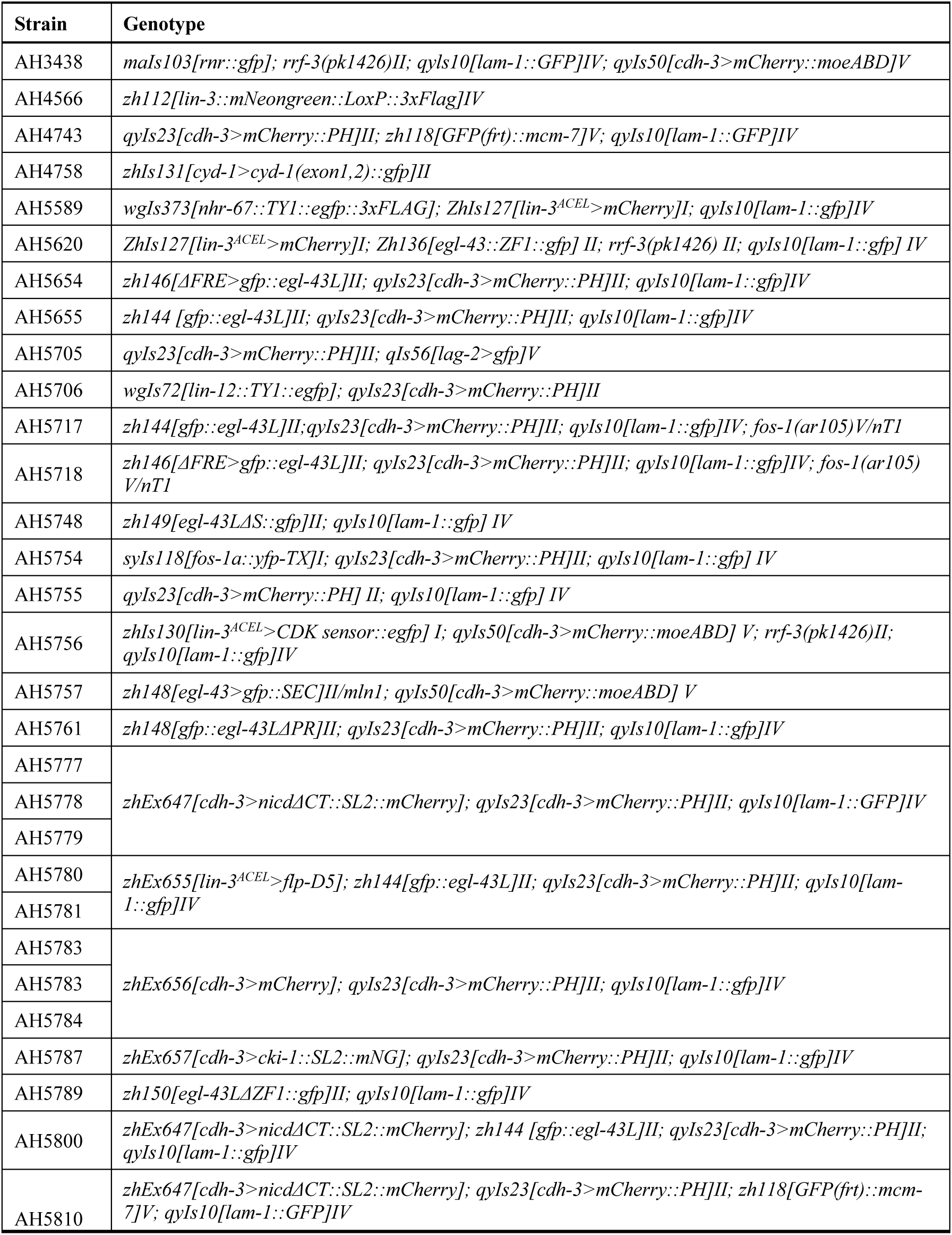
List of strains used.

**Table S2.**
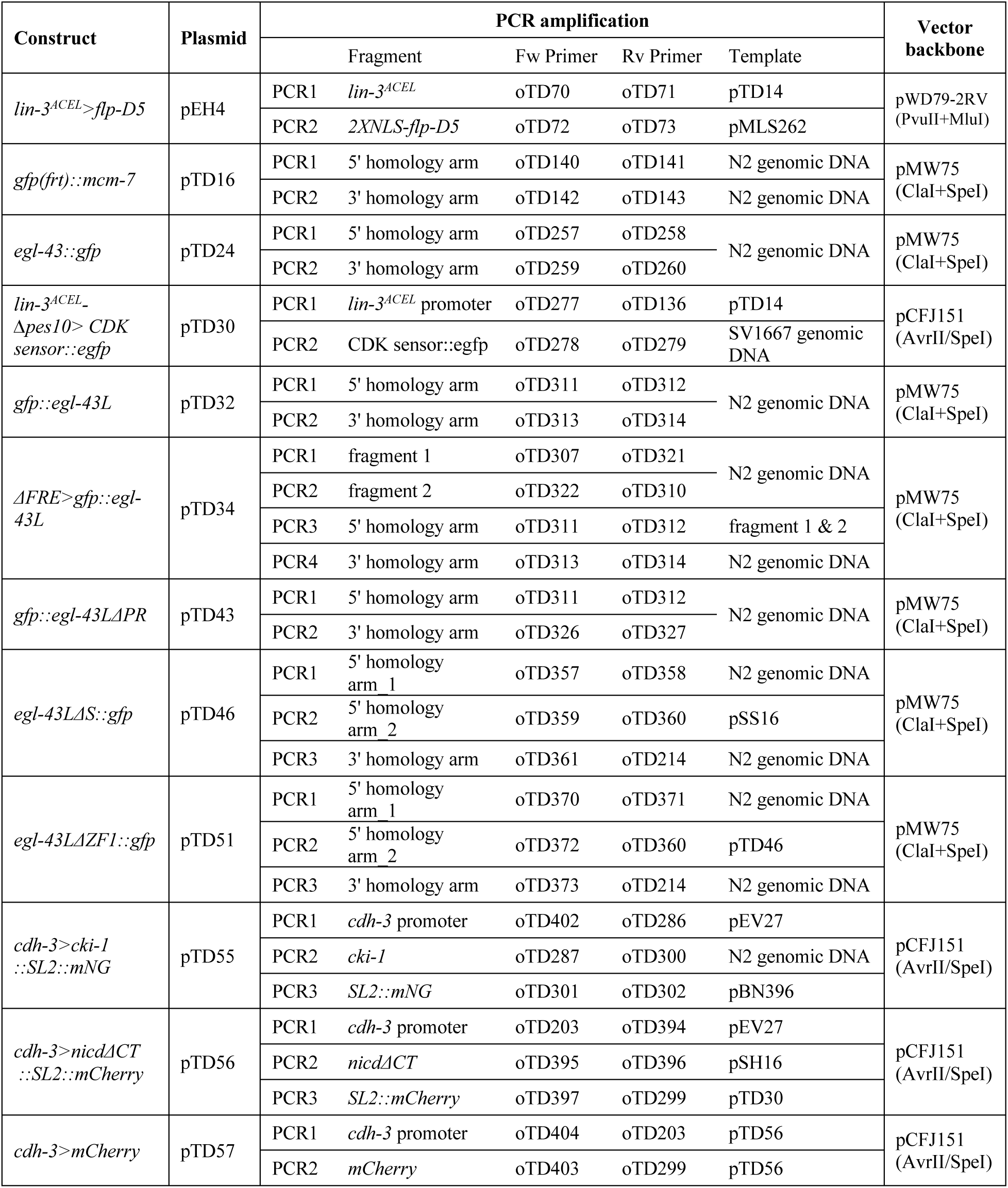
Design of plasmids used.

**Table S3.**
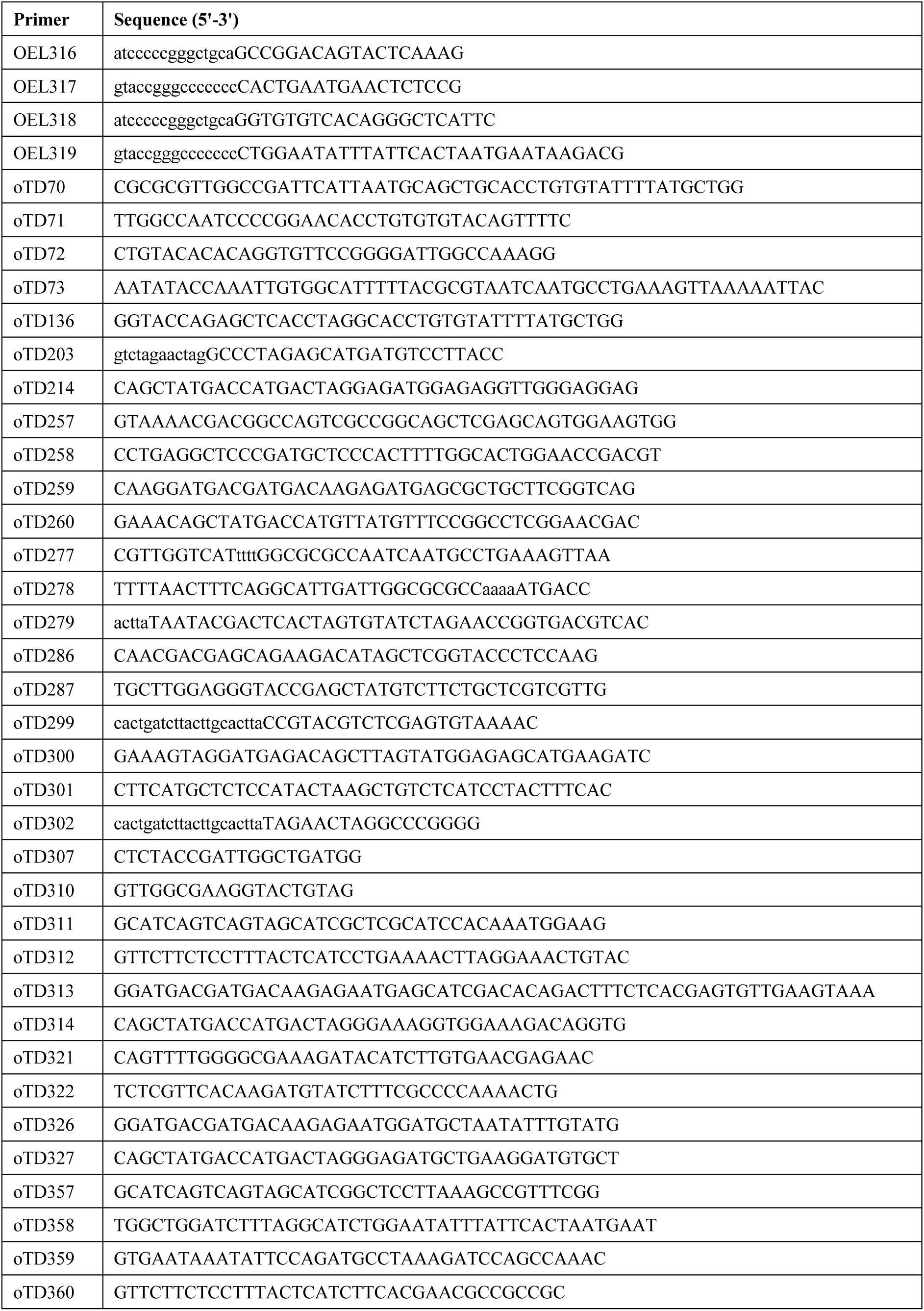
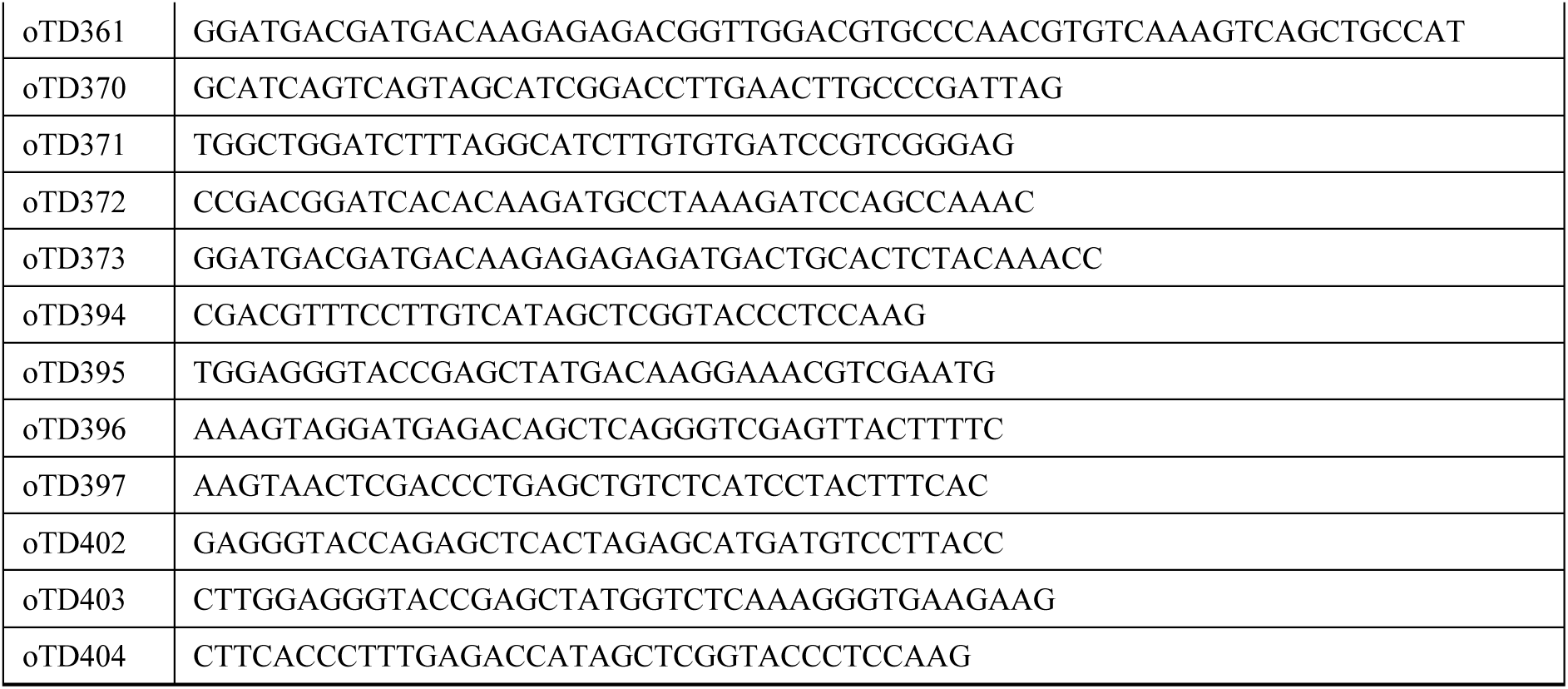
Oligonucleotide primers used.

**Table S4.**
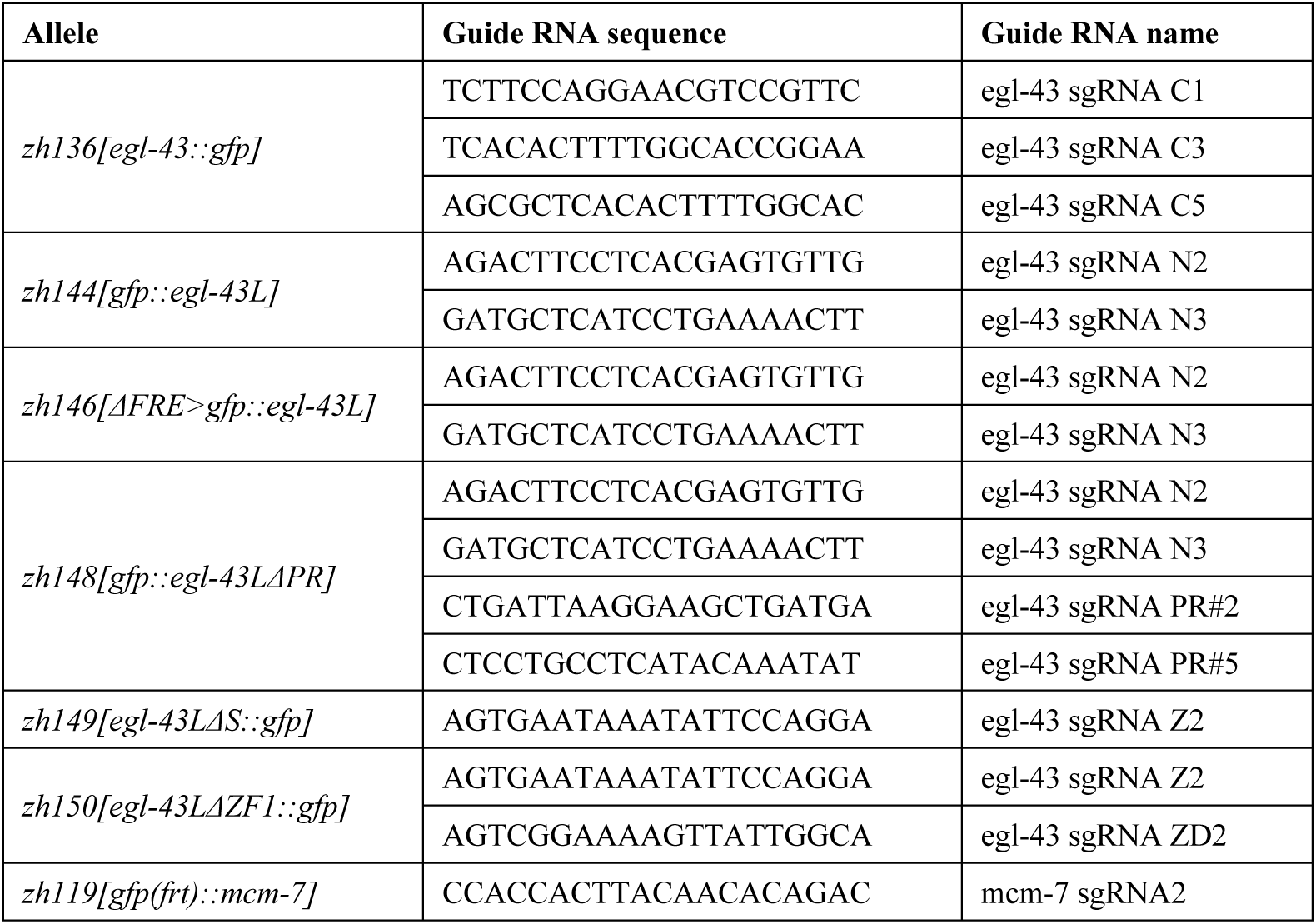
Sequences of the guide RNAs used.

### Extended methods

#### Generation of reporters and rescue transgenes

For all constructs, PCR fragments were amplified with Phusion DNA Polymerase (New England Biolabs) and were assembled by Gibson cloning (assembly kit #E2611, New England Biolabs). Details on the constructions of the plasmid and sequences of the oligonucleotides used can be found in suppl. Table S2 and S3. All plasmids were verified by DNA sequencing. For each transgenic reporter, single-copy transgene insertions were created according to the MosSCI protocol by microinjection of 50 ng/µl of the respective reporter plasmid together with 50 ng/µl pJL43.1 (*mos-1* transposase) and 2.5 ng/µl pCFJ90 (*myo-2>mCherry*), 5 µ pcFJ104 (*myo-3>mCherry*) and 10 ng/µl pGH8 (*rap-3>mCherry*) as co-injection markers as described [31].

#### Generation of endogenous reporters/deletions by CRISPR/Cas9

The CRISPR/Cas9 method described in [14] was used to generate the endogenous *egl-43* reporters and domain/motif deletions. For the ΔFRE reporter, a 11 bp (TTACTCATCTT) deletion was introduced into the repair template plasmid (pTD34). For the ΔPR and ΔZF1 domain deletions, two sets of guide RNAs each targeting one deletion breakpoint were used. A list of the sgRNAs used for each CRISPR/Cas9 allele can be found in suppl. Table S4.

